# Conflict between short-and long-term experiences affects visual perception by modulating sensory or motor response system: evidence from Bayesian inference models

**DOI:** 10.1101/2023.05.22.541675

**Authors:** Jing-Yi Wang, Xiu-Mei Gong, Qi Sun

## Abstract

Huge studies have explored the effects of short-and long-term experiences on visual perception, respectively. However, no study investigated whether and how the conflict between the two types of experiences affected our visual perception. To address this question, we adopted a task of estimating simulated self-motion directions (i.e., headings) from optic flow, in which a long-term experience – straight-forward motion is more often than lateral motion – plays an important role. The long-term experience is learned daily or encoded in our brains from birth. The heading directions in the experiment were selected from three different distributions, generating different conflicts between short-and long-term experiences. The results showed that both estimation errors and sizes of serial dependence of previously seen headings on current heading estimates varied when the experience conflict changed. Finally, we developed two Bayesian inference models, assuming that the experience conflict affected visual perception by influencing the sensory representation’s likelihood distribution or motor decision process. We found that both models captured participants’ estimation errors and serial dependences well in three distributions. In conclusion, the current study revealed the effects of the conflict between short-and long-term experiences on visual perception and preliminarily uncovered that Bayesian inference theory could explain the effects. Moreover, the study implied that the experience conflict affected visual perception by modulating our sensory or motor response systems.

## Introduction

Learning the statistical regularity of the living environment is vital for human and animal survival. For example, our ancestors discovered that red tomatoes are ripe and edible during the long evolutionary process. As a result, when shopping for tomatoes, we prefer fresh red tomatoes over green ones. Another example is that a person sees a hole in the road when driving to work. The person will slow down in advance when getting off work and passing by the same place. Hence, experiences have important effects on our daily behaviors.

According to the duration that the experience is stored, the experience can be categorized as long-term and short-term experiences. Massive studies have demonstrated that *the short-term experience, learned in the last few minutes or hours,* affects human perceptual behaviors. The most representative phenomenon is central tendency meaning that feature estimates are systematically compressed to the mean of the short-term experience about the feature (i.e., the distribution of learned features) [1]. For example, Jazayeri and Shadlen [2] asked participants to estimate the time interval in three experimental sessions. Time intervals were selected from three uniform distributions, but the shortest and longest times of each distribution were different, creating short, intermediate, and long distributions. Their results showed that time interval estimates were systematically compressed toward the distribution mean in each session, showing central tendency. Similar patterns have also been found for the perception of other features, such as line length [3-5], numerosity [6], and color [7, 8]. Taken together, these all demonstrate the effects of short-term experience on feature perception.

Note that in the studies above, short-term experience means the statistical regularity of features learned in the past few minutes or hours. To differentiate it, we define *long-term experience as the statistical regularity of features accumulated daily or in the long evolutionary process rather than learned in several days, which is robust and cannot be changed easily*. For example, there are more vertical and horizontal orientations than other directions in the natural world [9, 10], and straight-forward motion is more often than lateral motion. Studies have demonstrated that these long-term experiences affect our perception. For example, observers’ orientation estimates are systematically biased away from vertical and horizontal orientations, showing tilt or oblique illusion [11-14]; and observers’ self-motion direction (i.e., heading) estimates are systematically compressed toward the straight-ahead direction, showing center bias [15-19]. These all demonstrate the effects of long-term experience on feature perception. Levari et al. [20] asked participants to complete a color discrimination task (blue vs. purple). In the initial 200 trials, the blue and purple dots each accounted for 50%. In the last 200 trials, the proportion of blue dots were reduced to 6% in the decreasing condition or held constant in the stable condition. They found that participants tended to report more blue colors in the decreasing condition than in the stable condition. They named the phenomenon the “prevalence-induced concept change (PICC)” effect [21]. With their experimental methods, we know that the color distributions of the initial and last 200 trials were learned by participants within hours. Hence, the experiences are short-term, indicating that the PICC effect reflects the effects of conflicts between short-term experiences on perception, which led us to ask whether and how the conflict between short-and long-term experiences affected our perception.

Moreover, beyond the perceptual errors mentioned above (e.g., central tendency, oblique effect, center bias, PICC effect), researchers have found that our current perception is affected by a single feature that was previously seen several seconds earlier (within ∼15 s), known as serial dependence (see [22] for a review). For example, Fischer and Whitney [23] were the first to find that orientation estimates of current trials were systematically biased toward the orientations of previous trials. Similar result patterns are also observed in luminance [24], spatial location [25], motion direction [26], heading direction [16, 18, 27], numerosity [28], face identity [29], and so on. Serial dependence mainly indicates the effects of a past event on the current perception. Given that the past event is a component of the short-term experience, we raised another question of how serial dependence changed when short-and long-term experiences conflicted.

To sum up, the current study aimed to examine whether and how the conflict between short-and long-term experiences affected the perceptual estimate and serial dependence in visual perception. To address this question, we asked participants to estimate their heading direction from optic flow (Figure 1a and Movie 1 download from https://osf.io/4g5qk, [16-18]), in which the long-term experience affects its estimates [18, 19]. We selected heading directions simulated by optic flow from three different distributions (Figure 1b) to generate different conflicts between short-and long-term experiences. Our results showed that the heading estimate and serial dependence varied with the change in the experience conflicts. Finally, we developed two Bayesian observer models based on two assumptions: Model 1 assumed that the experience conflict affected the certainty of the sensory systems’ representation that corresponds the likelihood distribution in the Bayesian models. Model 2 assumed that the experience conflict affected participants’ motor decision systems. Specifically, the predicted estimates were scaled based on the level of the experience conflict. We found that the two models predicted participants well, implying that the experience conflict affected visual perception by modulating sensory or motor response systems. The current study improves our understanding of the effects of short-and long-term experiences on visual perception.

**Figure 1.**
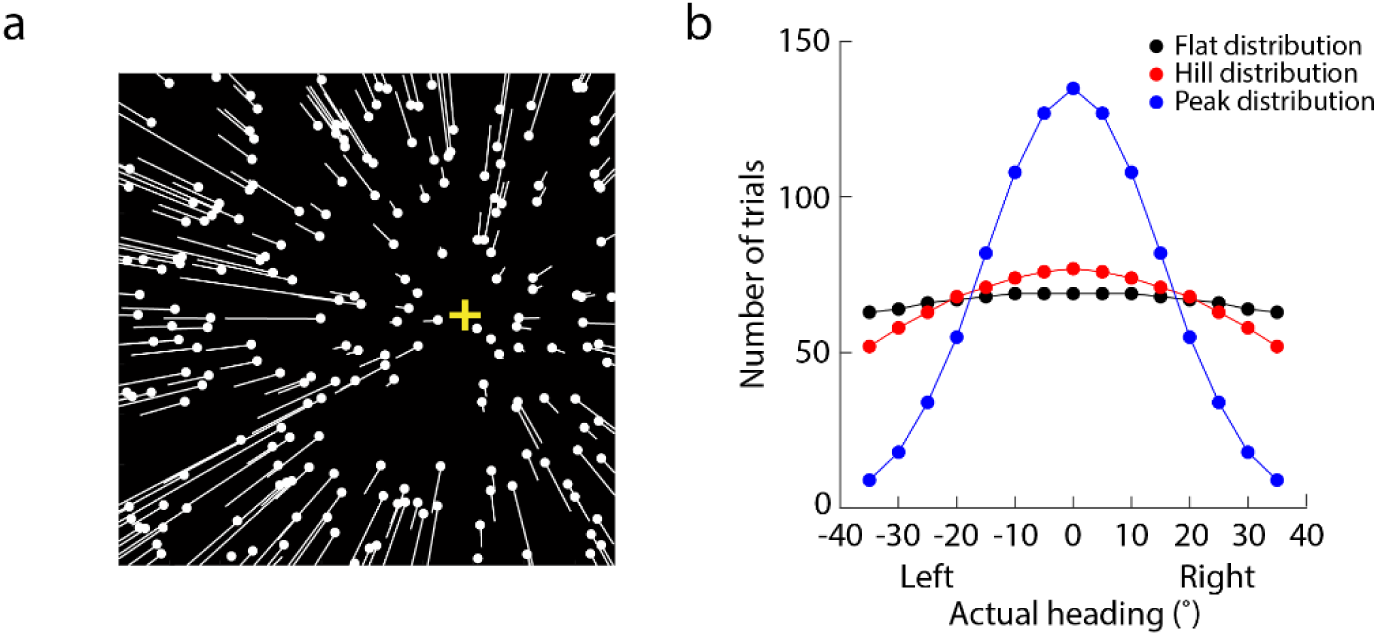
(a) Illustration of one optic flow display (see also Movie 1, https://osf.io/4g5qk). Optic flow displays simulated observers translating in a 3D dot-cloud that consisted of 126 dots. In the figure, the white dots indicate the dots’ initial positions in one trial. The white lines indicate the dots’ moving trajectories in the following frames. The longer the lines are, the faster the dots move. The yellow “+” indicates the simulated heading direction of the current optic flow. The lines and “+” are invisible in the experiment. (b) Number of trials plots against the actual heading. “Left” and “Right” on the x-axis represents that the actual heading is left or right to the straight-ahead direction (0 degrees). Black, red and blue dots correspond the flat, hill and peak distributions which are bell shaped with different widths and heights.

## Experiment

### Methods

#### Participants

Three groups of participants were enrolled. Each group had 12 participants (4 males, 8 females) and established one condition. All were naïve to the purpose of the experiment, and had a normal or corrected-to-normal vision. We obtained written informed consent forms from all participants before the commencement of the experiment. The consent form was approved by the Institutional Review Board at Zhejiang Normal University.

#### Visual stimuli and apparatus

In the experiment, participants were shown a serial of optic flow displays [30] (Figure 1a) that simulated observers translating at 1.53 m/s through a 3D random-dot cloud (80 degrees × 80 degrees, depth range: 0.20 m – 5 m). The 3D dot-cloud consisted of 126 dots (diameter: 0.21 degrees). The simulated directions of observers’ translation (headings) were ±35, ±30, ±25, ±20, ±15, ±10, ±5, and 0 degrees. Positive and negative values corresponded to heading to the right or left of the straight-ahead direction (0 degrees), respectively.

The stimuli were generated on a DELL workstation with an NVIDIA GeForce GTX 970 graphics card at a frame rate of 120 Hz and presented on a DELL monitor (112 degrees x 80 degrees) with a resolution of 2560 x 1440 pixels.

#### Experimental design and procedure

Participants conducted the experiment in a light-excluded room with their heads stabilized by a chinrest. They monocularly viewed the display with their dominant eye at a viewing distance of 20 cm, which aimed to reduce the conflicts between the motion parallax and binocular disparity. Participants’ viewing direction (i.e., straight-ahead direction) was aligned with the display center.

The current study consisted of three blocks, each completed by one group of participants, respectively. Each block corresponded to one distribution condition (Figure 1b): flat, hill, and peak distributions. The numbers of trials in the three distributions were generated from a Gaussian function with different standard deviations (σ = 80, 40, and 15 degrees) and the same mean (μ = 0 degrees). Therefore, the smaller the σ, the thinner and higher the distribution. Particularly, the peak distribution with a standard deviation of 15 degrees may be most similar to the long-term experience based on the tuning curve of the MSTd area in the monkey brain ([31, 32]). Therefore, the three distributions generated different conflicts between long-term and short-term experiences. The sense of conflict may be the weakest in the peak distribution, while the sense of conflict is strongest in the flat distribution. In the flat distribution (black dots in Figure 1b), the numbers of trials of ±35, ±30, ±25, ±20, ±15, ±10, ±5, and 0 degrees were 63, 64, 66, 67, 68, 69, 69, and 69. Hence, the number of trials were very similar and symmetric about the straight-ahead direction (0 degrees). In the hill distribution (red dots in Figure 1b), the numbers of trials of ±35, ±30, ±25, ±20, ±15, ±10, ±5, and 0 degrees were 52, 58, 63, 68, 71, 74, 76, and 77. Hence, the distribution had a low peak and looked like a hill. In the peak distribution (yellow dots in Figure 1b), the numbers of trials of ±35, ±30, ±25, ±20, ±15, ±10, ±5, and 0 degrees were 9, 18, 34, 55, 82, 108, 127, and 135. This distribution was thin and high. Each block was repeated two times, so there were a total of 2002 trials in each distribution.

The trial procedures of the three blocks were the same. On each trial, an optic flow display was presented for 500 ms, followed by a blank display with a horizontal line appearing across the mid-screen of the display. Participants were asked to indicate their perceived heading by moving a mouse-controlled probe. No feedback was given in the experiment.

Before the commencement of the experiment, participants were asked to conduct 20 practice trials randomly selected from the experimental distribution to get familiarized with the experiment. The experiment lasted for about 50 min.

### Data analysis

Firstly, previous studies have shown center bias in heading perception from optic flow [16-18]. Hence, we first examined whether and how the conflict between short-and long-term experiences affected center bias in heading perception. The data analysis methods were similar to that of Sun et al. [17]. Specifically, we fitted the perceived heading (PH) as a linear function of the actual heading (AH), given as:

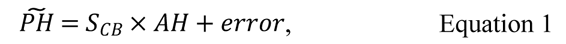

in which *PH* was the predicted perceived heading, *S*_*CB*_ was the slope. If center bias was presented in heading perception, then *S*_*CB*_ would be significantly smaller than 1. The larger the *S*_*CB*_, the samller the size of center bias. Importantly, a one-way repeated-measures ANOVA was conducted to examine whether the distributions affected the size of center bias. If the main effect of distributions was significant, then center bias was affected by heading distributions. According to the PICC effect meaning that participants tended to bias their estimates toward the features whose proportions were reduced in the short-term experience. We, therefore, expected that center biases in the flat, hill, and heavy-tail distributions were stronger than in the peak distribution.

Next, we tested how the conflict between short-and long-term experiences affected the serial dependence in heading perception. To evaluate serial dependence, we first calculated the heading error (HE) of each trial and the relative heading (RH). HE was the difference between the perceived and actual headings; RH was the difference in actual heading between the previous 1^st^ trial and current trial.

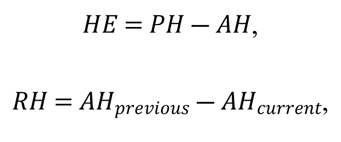

Then, the heading error (HE) was fitted as a linear function of the relative heading (RH), given as:

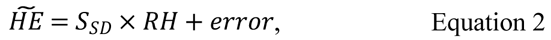

in which *HE* was the predicted heading error, *S*_*SD*_ was the slope. If serial dependence was presented in heading perception, then *S*_*SD*_ would be significantly different from 0. And, if *S*_*SD*_ was significantly less than 0, then a repulsive serial dependence was present in heading perception, meaning that the perceived heading of the current trial was biased away from the previously seen headings; conversely, if the *S*_*SD*_ was significantly larger than 0, then an attractive serial dependence was present in heading perception, meaning that the perceived heading of the current trial was biased toward the previously seen headings.

Note that the serial dependence analysis above did not remove the heading error induced by center bias from the total heading error. Next, we used the perceived heading (PH) minus the predicted perceived heading by Equation 1 (*PH*). The residual heading error (RHE) was proposed to be induced by the serial dependence after removing center bias [17]. Then, we fitted the residual heading error (RHE) as a function of the relative heading (RH). We adopted two types of functions based on the data trends in different distributions. For the data in the flat and hill distributions, we fitted the RHE as a 1^st^-derivative Gaussian function of RH (see [23]), given as:

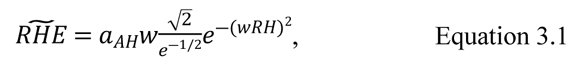

in which *RHE* was the predicted residual heading error, *a*_*AH*_ was the two amplitude of curve peaks, *w* was the curve width.

For the data in the peak distribution, we fitted the RHE as a linear function of the RH (see [17]), give as:

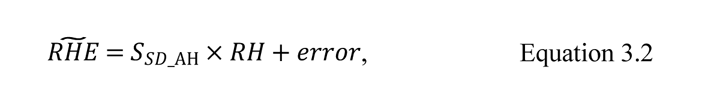

in which *S*_*AH_SD*_was the slope.

A bootstrapping test was conducted to test whether *a*_*AH*_ and *S*_*SD*_AH_ were significantly smaller or larger than 0. Specifically, we sampled the data of 12 participants 10,000 times with replacements. For each bootstrapping, we fitted the sampled data with Equations 3.1 or 3.2. The procedure generated a distribution of sampled *a*_*AH*_ and sampled *S*_*SD*_AH_. If the lower bound of the 95% CI of the distribution was larger than 0, then *a*_*AH*_ or *S*_*SD*_AH_ was significantly larger than 0, indicating an attractive serial dependence in heading perception; if the upper bound of the 95% CI of the distribution was smaller than 0, then *a*_*AH*_ or *S*_*SD*_AH_ was significantly smaller than 0, indicating a repulsive serial dependence in heading perception.

The relative heading (RH) in the above analysis was the difference in the actual heading between the previous and current trials, revealing a serial dependence in the sensory representation stage (i.e., perceptual stage, [23, 33, 34]). Some previous studies have also pointed out the motor decision process [35]. Next, to examine whether the motor decision process played some role in our serial dependence, we replaced the actual heading of previous trials with their perceived heading. The difference was named perceived relative heading (PRH). We then replaced the RH in Equations

### 3.1 and 3.2 with the PRH

To differentiate the serial dependences induced by the actual heading and the perceived heading of previous trials, *a*_*AH*_ in Equation 3.1 became *a*_*PH*_, *S*_*SD*_AH_ became *S*_*SD*_PH_. Similar bootstrapping analysis was adopted to examine whether *a*_*PH*_ and *S*_*SD*_PH_ were significantly smaller or larger than 0. In addition, if 95% CIs of *a*_*AH*_ and *a*_*PH*_ were overlapped, then the motor decision process did not induce extra serial dependence, so were the 95% CIs of *S*_*SD*_AH_ and *S*_*SD*_PH_.

## Results

### Center bias

Figure 2a shows the results of center bias in three distribution conditions. The x-axis (y-axis) was the actual (perceived) heading. The left panel shows the results of flat, hill and peak distributions. The right panel shows the results of peak and heavy-tail distributions. The solid lines and dashed blue line are the best fitting results of Equation 1. It clearly shows that all slopes were significantly smaller than 1 (*t*s (11) < 3.50, *p*s < 0.001, Cohen’s ds > 2.11), indicating center bias in heading perception.

**Figure 2.**
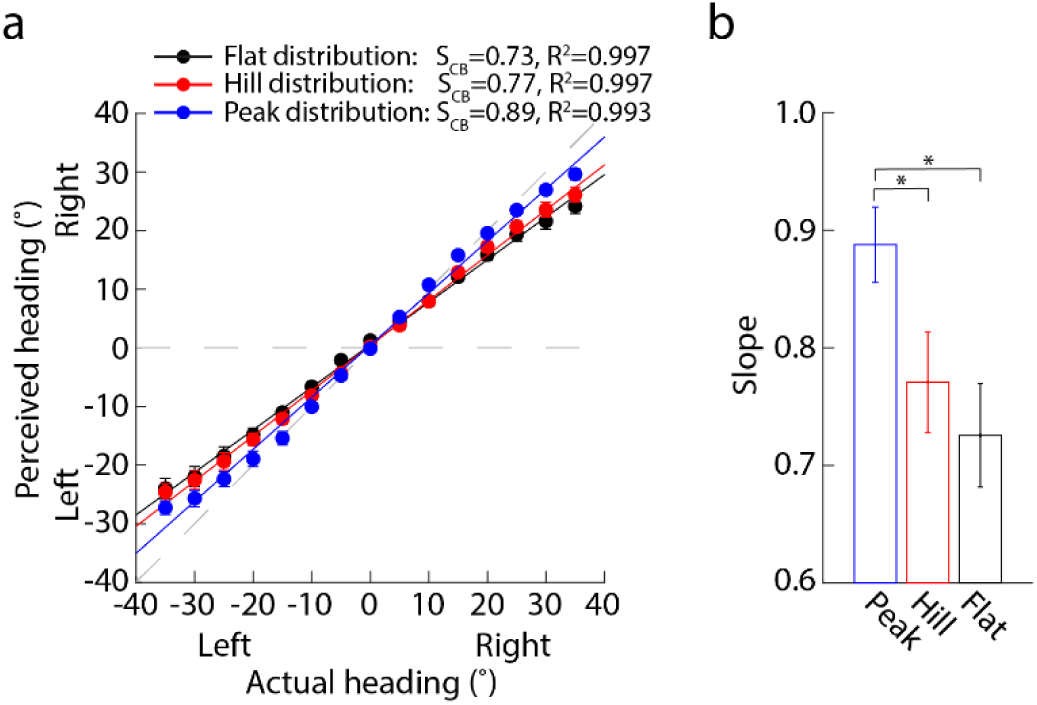
Center bias results. (a) x-axis (y-axis) is the actual (perceived) heading. Left and Right on the x-axis (y-axis) mean that the actual (perceived) heading is left or right to the straight-ahead direction (0 degrees). Black, red and blue dots correspond to the flat, hill and peak distributions. The horizontal gray dashed line indicates the pure center bias, meaning that perceived headings are always 0 degrees (i.e., straight-ahead direction) regardless of actual headings. The diagonal gray dashed line indicates the perfect performance, meaning that perceived headings are equal to actual headings. Each dot is the mean perceived heading averaged across 12 participants. Error bar is the standard error across 12 participants. (b) The slope (*S_CB_*) of Equation 1 is against four distributions. Error bar is the standard error across 12 participants. *, *p* < 0.05; **, *p* < 0.01; ***, *p* < 0.001.

Taking the peak distribution as the baseline (blue dots), Figure 2a clearly shows that when the proportions of peripheral headings in the hill and peak distributions (black and red dots, Figure 1b) increase, the slopes gradually decrease, suggesting that the sizes of center bias increase. One-way repeated-measures ANOVA analysis showed that the main effect of distributions was significant (*F* (2, 22) = 4.46, *p* = 0.024, η2 = 0.29). Further post-hoc analysis (Figure 2b) showed that the slopes of flat (Mean±SE: 0.73±0.044) and hill (0.77±0.043) distributions were all significantly smaller than that of the peak distribution (0.89±0.032) (*p*s < 0.049), indicating that the size of center bias increased when the proportions of peripheral headings increased in the short-term experience.

### Serial dependence

Figure 3a plots the heading error against the relative heading (the difference in the actual heading between the previous 1^st^ trial and current trial), showing the serial dependence effect before removing center bias. Three panels clearly show that when the actual heading of the previous 1^st^ trial is left (right) to the actual heading of the current trial, the heading estimate of the current trial was left (right) to the actual heading, showing an attractive serial dependence. One sample *t*-test showed that the slopes of three distributions were all significantly larger than 0 (*t*s (11) > 2.38, *p*s < 0.037, Cohen’s ds > 1.43). One-way repeated-measures ANOVA showed that the main effect of distributions was significant (*F* (2, 22) = 4.63, *p* = 0.021, η2 = 0.30). Further post-hoc analysis (Figure 3b) showed that the slopes of flat (Mean±SE: 0.15±0.022) and hill (0.12±0.025) distributions were all significantly larger than that of the peak distribution (0.053±0.022) (*p*s < 0.044), indicating that before center bias was removed, the size of attractive serial dependence increased with increasing proportion of peripheral headings in the short-term experience.

**Figure 3.**
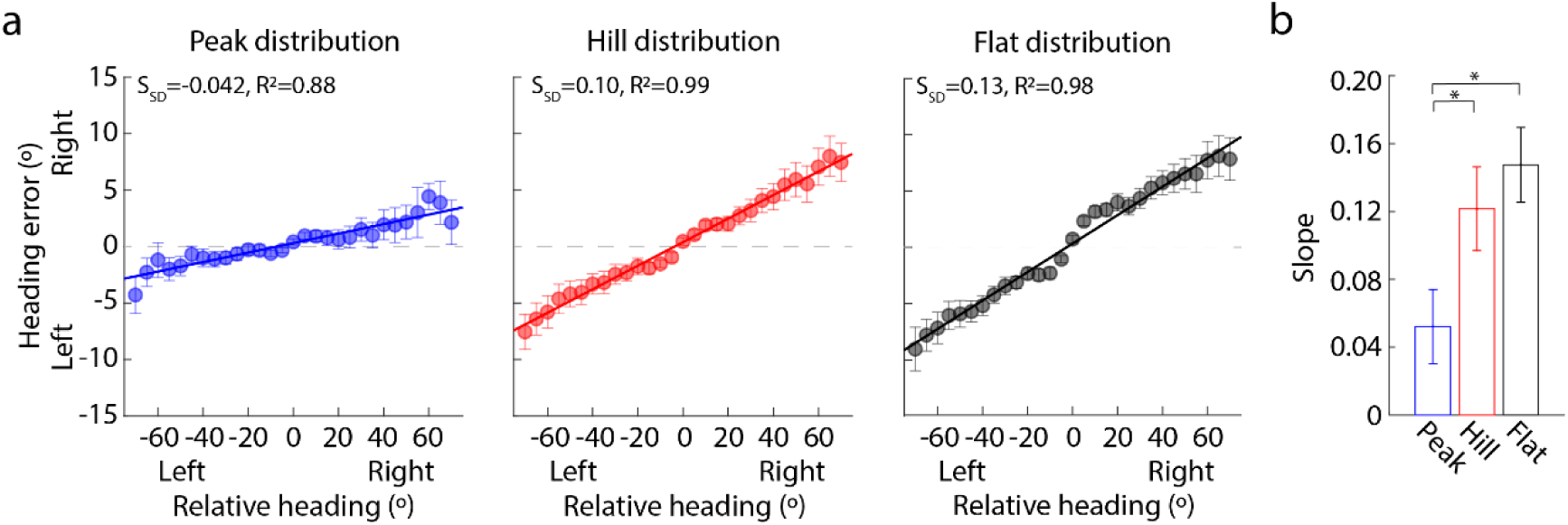
Serial dependence before removing center bias. (a) Heading error is against the relative heading in three distributions (e.g., flat, hill, and peak distributions). Heading error was the difference between perceived heading and actual heading. Relative heading was the difference in the actual heading between the previous 1^st^ trial and current trial. Each line shows the best fitting result of Equation 2. Each dot is the mean heading error averaged across 12 participants. Error bar is the standard error across 12 participants. (b) Slope is against the three distributions. Error bar was the standard deviations across 12 participants. *, *p* < 0.05; **, *p*<0.01; ***, *p*<0.001.

Next, we calculated the residual heading error by subtracting the error induced by center bias from the total heading error. Then, we fitted the residual heading error as a function of the relative heading (Equations 3.1 and 3.2). Figure 4a shows that after center bias was removed, in the peak distribution condition, the heading estimate of the current trial tended to be left (right) to the actual heading of the current trial when the actual heading of previous 1^st^ trial was right (left) to the actual heading of the current trial, showing a repulsive serial dependence (Bootstrapping test, 95% CI of *S*_*SD*_AH_ [-0.023 -0.0009], left panel in Figure 4b); however when there were more peripheral headings in the distributions, the serial dependence became attractive (hill: 95% CI of *a*_*AH*_ [0.42 0.95]; flat: 95% CI of *a*_*AH*_ [1.32 1.93], middle and right panels in Figure 4b). Importantly, the size of attractive serial dependence increased with increasing the proportion of peripheral headings in the short-term experience.

**Figure 4.**
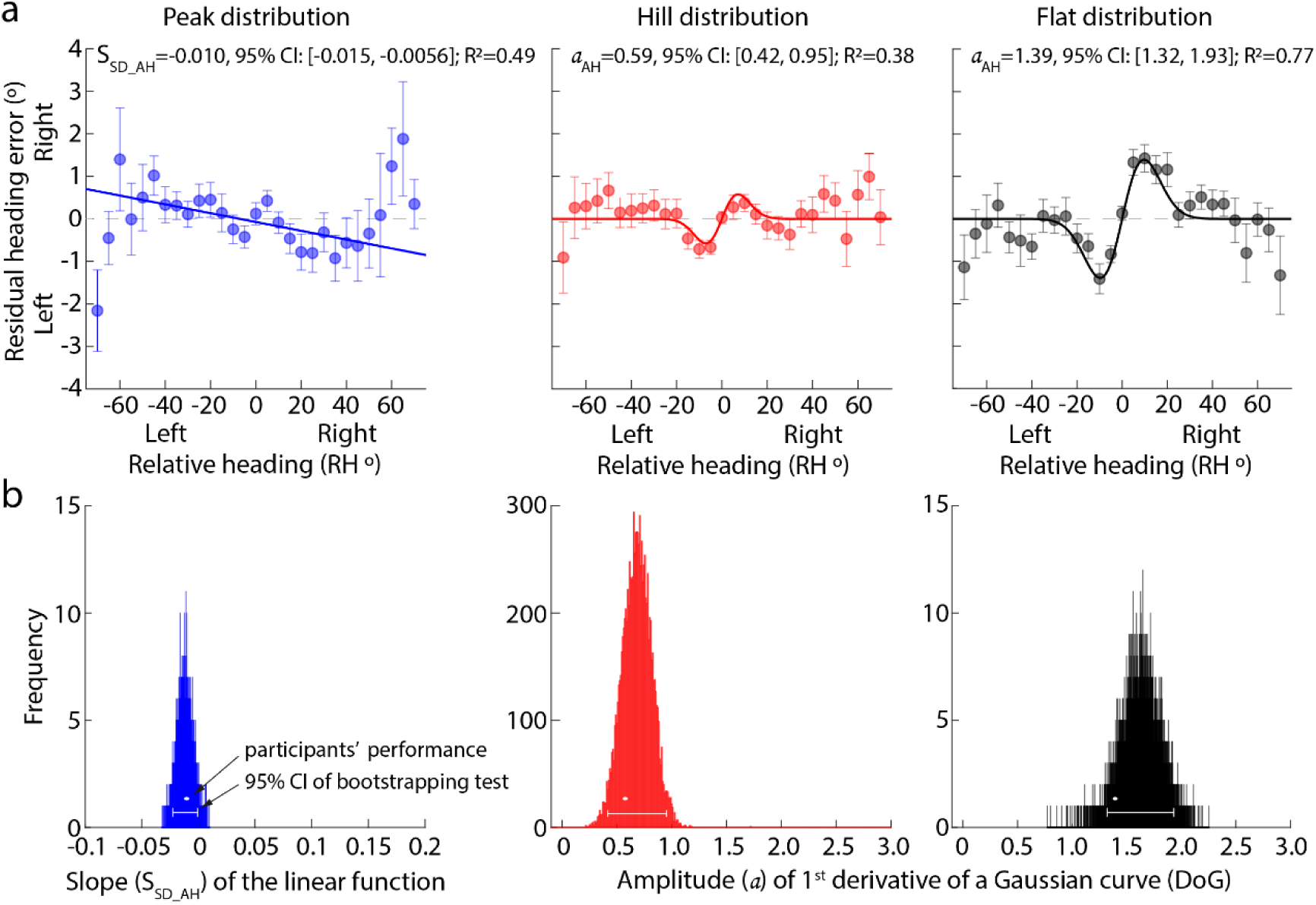
Serial dependence of the previous actual heading after removing center bias. (a) Residual heading error is against the relative heading in three distributions (e.g., flat, hill, and peak distributions). Residual heading error is the difference between perceived heading and predicted perceived heading by Equation 1. Relative heading was the difference in the actual heading between the previous 1^st^ trial and the current trial. Each line shows the best fitting result of Equation 3.1 or 3.2. Each dot was the mean heading error averaged across 12 participants. Error bar was the standard error across 12 participants. (b) Histogram of the slope (S_SD_AH_) or the amplitude (a_AH_) with Bootstrapping for 10,000 times. The white dot indicates the participants’ estimate. Horizontal line indicates the 95% CI of the slope or amplitude.

Lastly, we calculated the relative heading between the heading estimate of previous 1^st^ trial and the actual heading of current trial to examine whether the motor decision process affected serial dependence in heading perception. Figure 5 shows the same trend under different distribution conditions. Specifically, a repulsive serial dependence (95% CI: [-0.023 -0.0056], left panels in Figure 5) was in the peak distribution condition; an attractive serial dependence was in the other distributions (hill, 95% CI: [0.57 1.29]; flat, 95% CI: [1.074 1.82], middle and right panels in Figure 5). In addition, the size of attractive serial dependence increased with the proportion of peripheral heading in the distribution. However, the sizes of serial dependence were not significantly different between the RH and PRH due the overlap of the 95% CIs. Therefore, the motor response process did not affect serial dependence in heading perception from optic flow.

**Figure 5.**
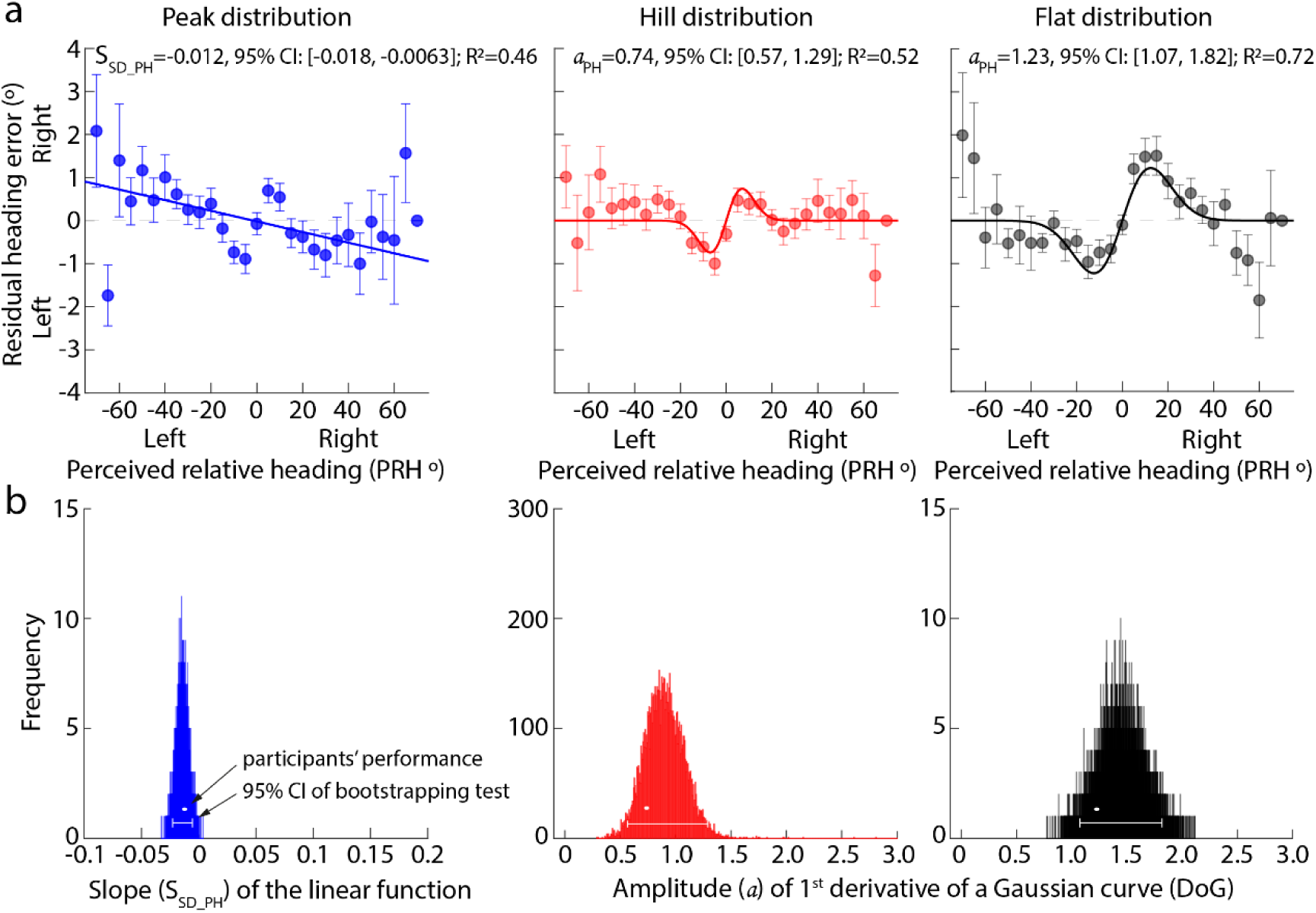
Serial dependence of the previous estimate after removing center bias. (a) Residual heading error against the relative heading in three distributions (e.g., flat, hill, heavy-tail, peak distributions). Residual heading error was the difference between perceived heading and predicted perceived heading by Equation 1. Relative heading was the difference in the actual heading between the previous nth trial and current trial. Each line shows the best fitting result of Equation 3.1 or 3.2. Each dot is the mean heading error averaged across 12 participants. Error bar is the standard error across 12 participants. (b) Histogram of the slope (S_SD_PH_) or the amplitude (a_PH_) with Bootstrapping for 10,000 times. The white dot indicates the participants’ estimate. Horizontal line indicates the 95% CI of the slope or amplitude.

## Discussion

To sum up, aside from reproducing center bias and serial dependence in heading perception from optic flow (e.g., [16-18, 27]), the current experiment confirmed that the conflict between short-and long-term experiences affected center bias and serial dependence in heading perception from optic flow.

Specifically, when there were more peripheral headings in the short-term experience than in the long-term experience, observers tended to bias their estimates away from these headings, leading to an increase in center bias in the hill and flat distributions (Figure 2). This finding is consistent with the prediction of the PICC effect [20, 21]. Unlike Levari’s PICC effect, which was due to the conflict between short-term experiences, we found that the PICC effect could also occur when short-and long-term experiences conflicted.

Moreover, the conflict between short-and long-term experiences could affect the serial dependence. When the headings were selected from the peak distribution, a repulsive serial dependence was in heading perception, reproducing the finding of Sun et al. [17], in which the headings were arranged in a geometric sequence (±32°, ±16°, ±8°, ±4°, ±2° and 0°), distributed like a peak distribution. When the headings were selected from a distribution that was similar to uniform distribution (e.g., flat), the serial dependence became attractive, reproducing the finding of Xu et al. [18], in which the headings (±30°, ±25°, ±20°, ±15°, ±10°, ±5°, and 0°) were uniformly distributed.

Lastly, the current study again found that the response decision process did not affect the serial dependence in heading perception, supporting the previous studies (e.g., [17, 18]).

### Bayesian inference model

Previous studies have proposed that heading perception from optic flow is a Bayesian inference process, meaning that the heading estimate is a result of an optimal combination between prior and likelihood distribution (gray shaded area in Figure 6, [17-19]). In the current study, we developed two Bayesian inference models based on two assumptions to explain the effects of the conflict between short-and long-term experiences on heading perception from two perspectives.

**Figure 6.**
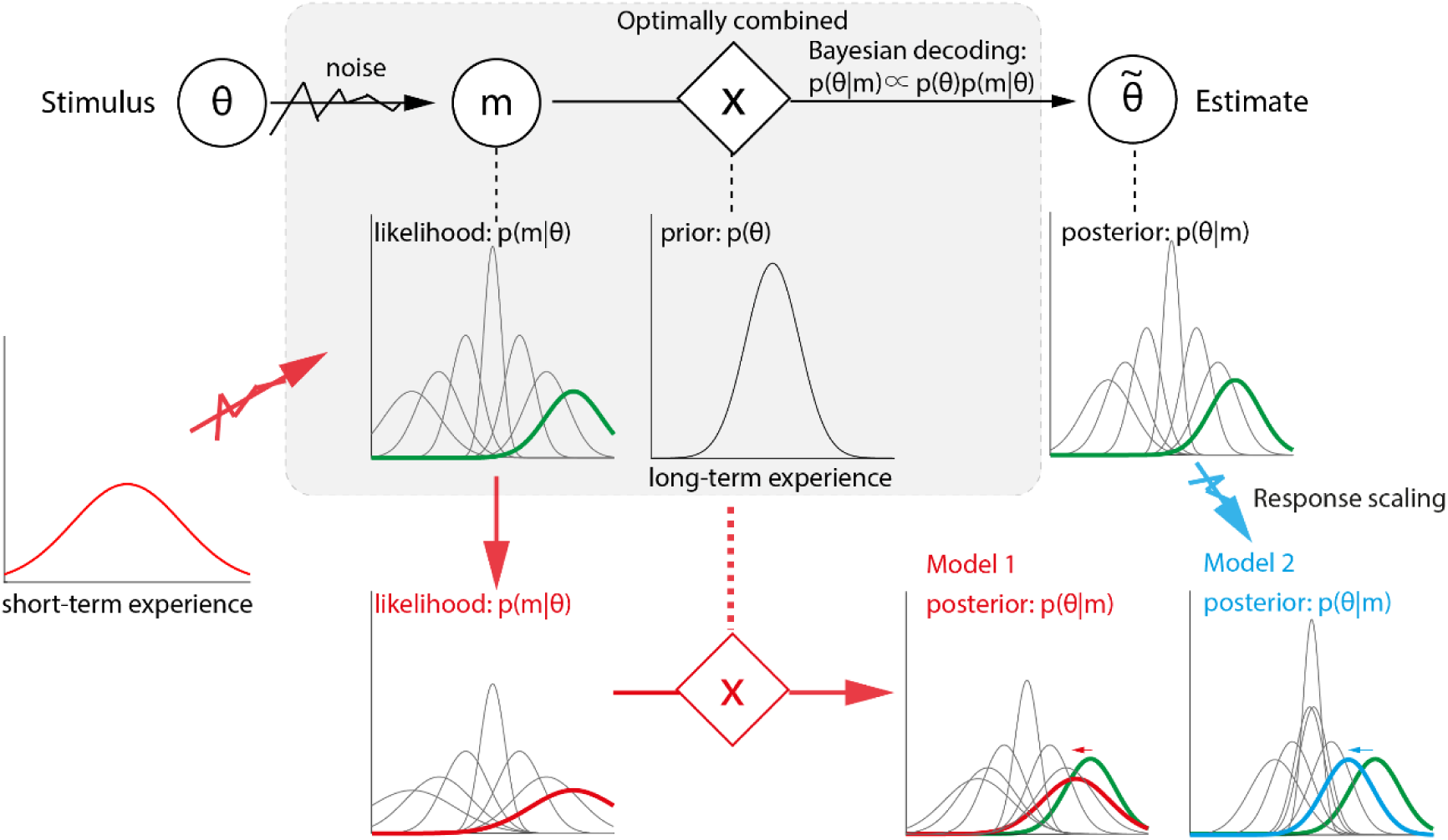
Illustration of Bayesian models. The gray shaded area illustrates the logic of the classical Bayesian inference model which proposes that the perceptual estimate is from a posterior distribution (p(θ|m)). The posterior is the result of an optimal combination of prior (p(θ)) and likelihood distribution (p(m|θ)). The red arrows illustrate our model 1 which proposes the short-term experience changes our likelihood distribution. For example, when there were peripheral headings, the sensory systems become fatigued. As a result, the certainty of the sensory representation decreases. The likelihood distributions become wider, as shown by the red line. The fatigued likelihood distribution makes the estimate more compressed toward the prior center, showing stronger center bias. Blue arrow illustrates our model 2, in which the likelihood distributions are the same as in the classical Bayesian inference model. However, when participants report by moving the mouse-controlled probe, the decoded estimates will be multiplied with a responding scaling factor, generating the final estimate from a shifted posterior distribution (e.g., blue line).

### Model 1 Conflict affects likelihood distributions

As mentioned above, the classical Bayesian inference process is an optimal combination of prior (*p*(*θ*)) and likelihood (*p*(*mθ*)) distributions (gray shaded area in Figure 6):

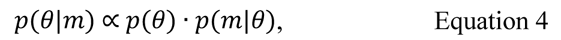

in which *θ* is the actual heading, *m* is participants’ estimate. *p*(*θ*|*m*) is the posterior distribution.

In the current study, the prior (*p*(*θ*)) was the long-term experience, following a Gaussian distribution with the straight-ahead direction (0°) serving as the distribution center, given by:

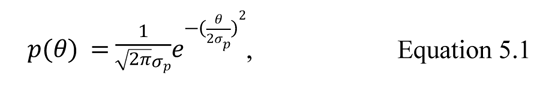

in which *σ*_*p*_ is a free parameter, indicating the standard deviation of the prior distribution. The smaller the *σ*_*p*_ is, the narrower the prior distribution is and the more certainty the prior is. In the model, we assumed that the prior distribution was robust and was not affected by the short-term experience.

In addition, the likelihood (*p*(*mθ*)) is also a Gaussian distribution with the actual heading of current trial (*θ*) serving as the distribution center, given by:

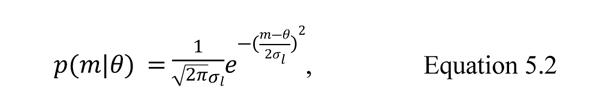

in which *σ*_*l*_ is another free parameter, indicating the standard deviation of the likelihood distribution. Like the *σ*_*p*_, the smaller the *σ*_*l*_ is, the narrower the likelihood distribution is and the more certainty the sensory representation of the current heading is.

Previous studies have found that when observers are often exposed to one feature (seconds to minutes), neurons of sensory systems selectively responding to the feature would become fatigued. Neurons’ firing rates are significantly reduced [36, 37]. Many studies have proposed that the likelihood distributions are closely related to neurons’ tuning curves [38]. Therefore, when the neurons became fatigued, the *σ*_*l*_ of the likelihood distribution was increased. Accordingly, there were more peripheral headings in the flat and hill distributions than in the peak distribution, leading to larger *σ*_*l*_ of these peripheral headings’ likelihood distributions in the flat and hill distribution than in the peak distribution; vice versa.

According to the Bayesian inference theory [39], estimates would be systematically compressed toward the center of the prior distribution (i.e., straight-ahead direction) with decreasing the certainty of likelihood distribution, showing an increase in center bias. Therefore, we expected that Model 1 could explain our current findings.

## Methods

As mentioned in the data analysis part of Experiment, the peak distribution severed as the baseline because we proposed that the peak distribution was most closely to our long-term experience. Therefore, we first built a classic Bayesian inference model (gray shaded area in Figure 6) to figure out the prior distribution and the likelihood distributions of different headings. Note that (1) we assumed that the standard deviations of two symmetric headings (e.g., ±35°) were the same; (2) and according to the previous studies [40], the standard deviation of the likelihood distribution increased with the heading value; (3) to improve the modeling efficiency and simplify the model, we assumed that the standard deviation of likelihood distribution linearly increased with the actual heading, given by:

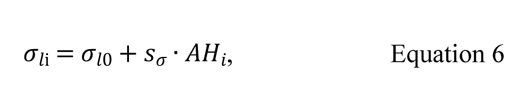

in which *σ*_*l*0_ is the standard deviation when the actual heading (AH) is 0°; *S*_*r*_ is the increasing factor and must be positive. Therefore, the model included three parameters: *θ*_*p*_, *θ*_*0*_, and *S*_*r*_.

Markov chain Monte Carlo (MCMC) sampling was used to estimate the parameters. We used a 1,000,000 iterations as a burn-in period and randomly select start-point values for the three parameters, e.g., [1 1 0.05]. The start-point values do not affect the following iterations. We first used the start-point parameters to fit participants’ data and calculated the sum of the log-likelihood posterior (LLP). From the second iteration, a random number selected from a standard norm distribution was added to each parameter of the pre-iteration. Then, we used the new parameters to fit participants’ data. We compared the sums of the LLPs between the current and previous 1^st^ iterations. If the sum of the LLP of current iteration is larger than that that of previous 1^st^ iteration, then the new parameters were kept and used in the next iteration. If not, a new random number (ε) was generated. If ε > 0.5, then the new parameters were kept; if ε ≤ 0.5, the parameters of previous 1^st^ iteration were kept. The same procedure was repeated for the left iterations. The final values of *σ*_*p*_, *σ*_*10*_, and *S* were the them mean of the 18,000 iterations selected from the 100,001st iteration to the end with step 50.

Next, we repeated the above procedure for the data in the hill and flat distribution. But, the *θ*_*p*_ of prior (*p*(*θ*)) was fixed and from the model of the peak distribution. Therefore, the models in the hill and flat distributions only contained two parameters: *σ*_*l*0_, and *S* (red arrows in Figure 6).

Finally, we used the ideal models to simulate 12 participants’ estimates in each actual heading under each distribution condition. Then, we sorted the sequence of the simulated trials according to the true participants’ trial sequences. Finally, we used the data analysis methods to analyze the simulated data. If the trends of the simulated center bias and serial dependence in different distributions were similar to participants’ center bias and serial dependence, then the effects of the conflict between short-and long-term distributions on heading perception were a Bayesian inference process. Meanwhile, the simulated results also suggested that the conflict between short-and long-term experiences affected heading perception by influencing the likelihood distribution of our sensory systems.

## Results and discussion

Figure 7 shows the simulated results of Model 1. Figure 7a plots the simulated perceived heading against the actual heading. It clearly shows that the pattern of simulated results was similar to that participants’ pattern (Figure 2a). Specifically, all simulated perceived headings were compressed toward the straight-ahead direction (horizontal dashed line), showing center bias; and the size of center bias increased with increasing the proportion of peripheral headings (peak to hill to flat distributions).

**Figure 7.**
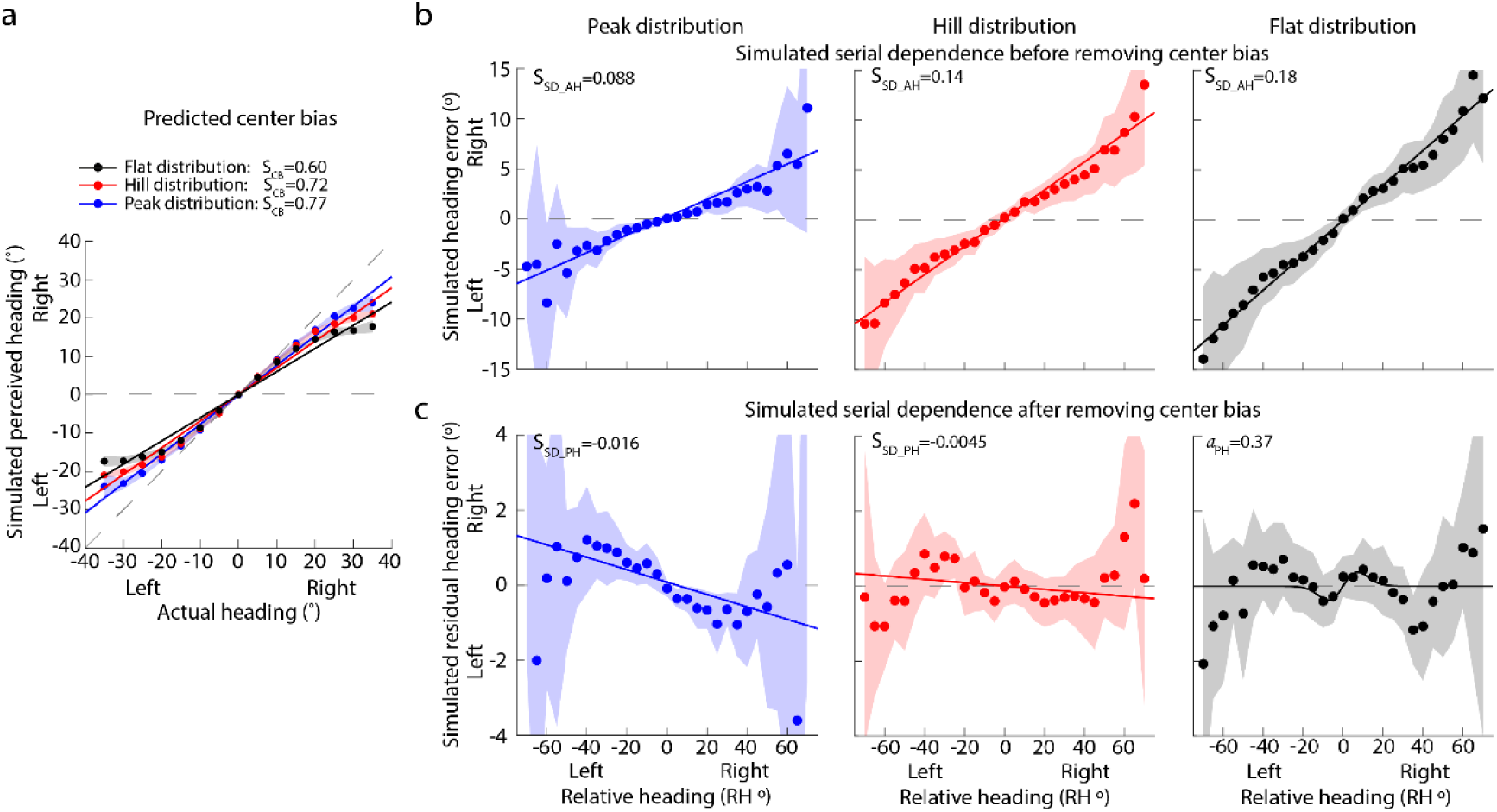
Simulated results of Model 1. (a) Simulated perceived heading against the actual heading. Left and Right on the x-axis (y-axis) mean that the actual (simulated perceived) heading is left or right to the straight-ahead direction (0 degrees). Black, red and blue dots correspond to the flat, hill and peak distributions. The horizontal gray dashed line indicates the pure center bias, meaning that perceived headings are always 0 degrees (i.e., straight-ahead direction) regardless of actual headings. The diagonal gray dashed line indicates the perfect performance, meaning that perceived headings are equal to actual headings. Each dot is the mean the simulated perceived heading averaged across 12 simulated participants. Error bar is the standard deviation across 12 simulated participants. (b) Simulated serial dependence before removing center bias. X-axis is the simulated heading error; y-axis is the relative heading – the difference in actual heading between the previous 1^st^ trial and the current trial. Each dot indicates the mean of the simulated heading error across 12 simulated participants in each relative heading. Shaded error means the standard deviation of the simulated heading error across 12 simulated participants. (c) Simulated serial dependence after removing center bias. Y-axis is the simulated residual heading error – the difference between the actual simulated perceived heading and the simulated perceived heading induced by center bias.

Figure 7b plots the simulated heading error against the relative heading, showing the serial dependence before removing center bias. It shows an attractive serial dependence and the size of serial dependence increased with the increasing the proportion of peripheral headings.

Figure 7c plots the simulated residual heading error against the relative heading, showing the serial dependence after removing center bias. It shows a repulsive serial dependence in the peak distribution and an attractive serial dependence in the flat distribution. The serial dependence was weak in the hill distribution.

Together, the patterns of our Model 1’s simulated results were similar to participants’ patterns (Figures 3 and 4), suggesting that the conflict between short-and long-term experiences may modulate the certainty of our sensory systems’ representation to affect heading perception from optic flow.

### Model 2 Conflict affects responses

Model 1 assumed that different performances in different distributions were due to the change of likelihood distributions. Some researchers could argue another explanation. In daily life, straight-forward motion is more often than lateral motion, leading observers to be reluctant to report the peripheral headings. The conflict between the short-and long-term experiences increased the sense of reluctance, leading to participants bias their estimates toward the center of the prior distribution. That is, the conflict between the short-and long-term experiences affected participants’ motor decision system rather than the likelihood distribution.

## Methods

Like Model 1, we also took the peak distribution as the baseline. The predicted performance can be given by Equation 4. The modeling procedure was the same as Model 1.

After getting the model for the peak distribution, we multiplied a response scaling factor (*r*) with the posterior estimates (*p*(*θ*|*m*)), as indicated by the blue arrows in Figure 6. Response scaling factor (*r*) can be given by:

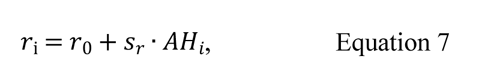

in which *r*_0_ is the standard deviation when the actual heading (AH) is 0°; *S*_*r*_ is the increasing factor and must be positive. Therefore, Model 2 contained two parameters: *r*_0_ and *S*_*r*_.Similar MCMC procedures were adopted to get the response scaling factors in the hill and flat distributions.

Lastly, similar procedures were adopted to predict the center bias and serial dependence in the flat and hill distributions (as see the last paragraph in the methods of Model 1).

## Results and discussion

Figure 8 shows the simulated results of our Model 2. It clearly shows that the center bias increases with increasing the proportion of peripheral headings (peak to hill to flat distributions); before removing center bias, the serial dependence was attractive in three distributions; and after removing center bias, a repulsive serial dependence was in the peak distribution and a very weak attractive serial dependence was in the flat distribution. The serial dependence was not evident in the hill distribution. Overall, the predicted result patterns of Model 2 were similar to participants’ patterns, suggesting that the conflict between short-and long-term experiences affects heading perception by modulating our motor decision system.

**Figure 8.**
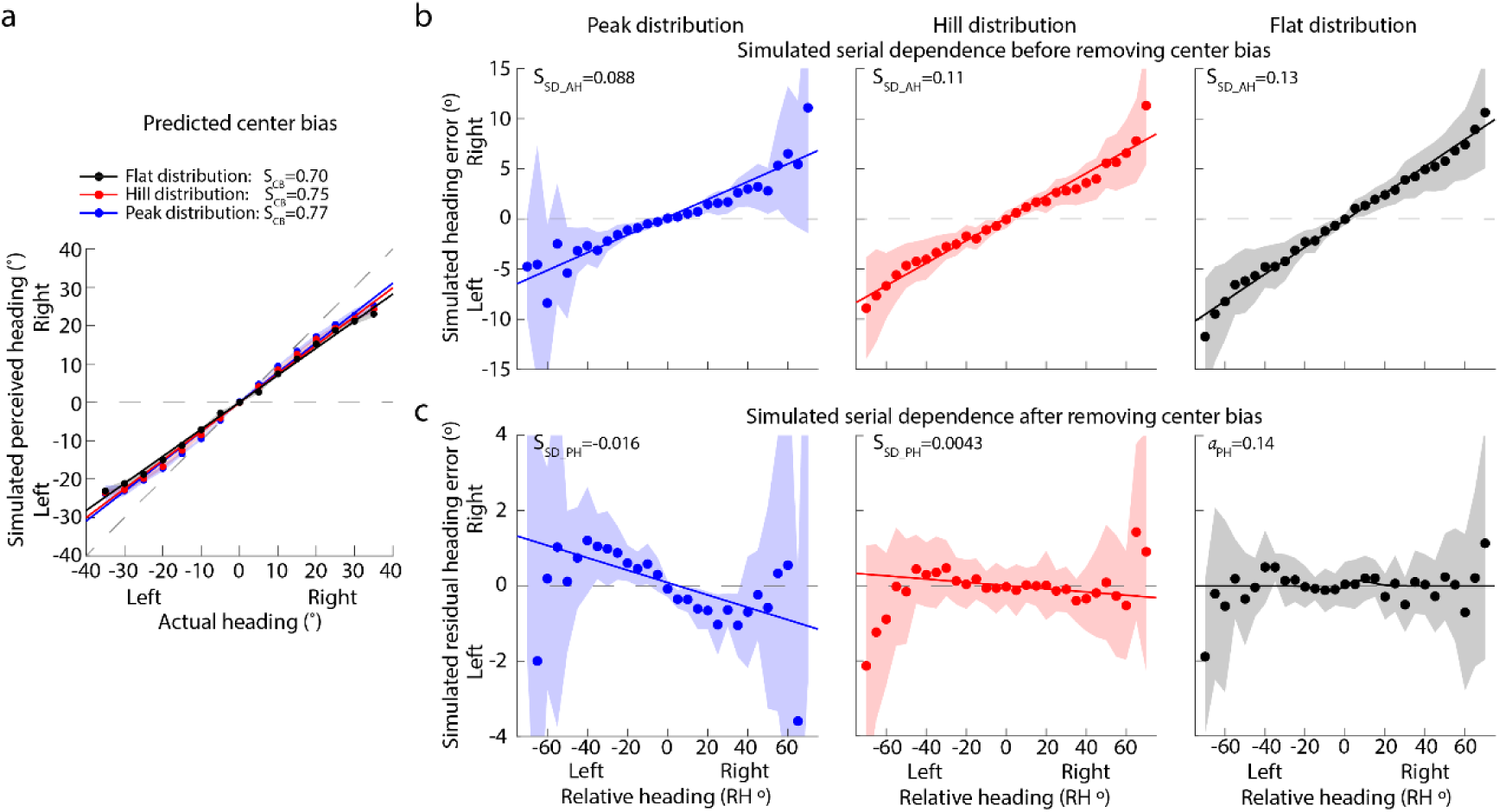
Simulated results of Model2. (a) Simulated perceived heading against the actual heading. Left and Right on the x-axis (y-axis) mean that the actual (simulated perceived) heading is left or right to the straight-ahead direction (0 degrees). Black, red and blue dots correspond to the flat, hill and peak distributions. The horizontal gray dashed line indicates the pure center bias, meaning that perceived headings are always 0 degrees (i.e., straight-ahead direction) regardless of actual headings. The diagonal gray dashed line indicates the perfect performance, meaning that perceived headings are equal to actual headings. Each dot is the mean the simulated perceived heading averaged across 12 simulated participants. Error bar is the standard deviation across 12 simulated participants. (b) Simulated serial dependence before removing center bias. X-axis is the simulated heading error; y-axis is the relative heading – the difference in actual heading between the previous 1^st^ trial and the current trial. Each dot indicates the mean of the simulated heading error across 12 simulated participants in each relative heading. Shaded error means the standard deviation of the simulated heading error across 12 simulated participants. (c) Simulated serial dependence after removing center bias. Y-axis is the simulated residual heading error – the difference between the actual simulated perceived heading and the simulated perceived heading induced by center bias.

By comparing the simulated results of the two models, we found that the simulated results of Model 2 were closer to the participants’ performance. The slopes of the simulated center bias were close to the slopes of the participants’ center bias, and so was the serial dependence before removing the center bias. However, in the flat distribution, Model 2’s attractive serial dependence was not stronger than Model 1’s. That is, Model 1 can better explain attractive serial dependence than Model 2. These made us propose that the effects of experience conflict could be due to the change in sensory systems and motor decision systems.

## General discussion

The current study adopted one behavioral experiment and two Bayesian inference models to systematically investigated whether and how the conflict between short-and long-term experiences affected the estimation error and serial dependence in visual perception. Previous studies have examined their independent effect on visual perception. Our results showed that the experience conflict affected estimation error and serial dependence in visual perception. Compared with the long-term experience in which peripheral headings were few, participants would estimate their headings with more errors and relied more on the previously seen heading to estimate their current heading when the short-term experience contained more peripheral headings, showing stronger center bias and attractive serial dependence. Meanwhile, our Bayesian inference models predicted participants’ performance well, suggesting that the effects of the experience conflict could be a Bayesian inference process. Importantly, we proposed that the experience conflict affected visual perception by modulating the certainty of sensory systems’ representation (e.g., likelihood distribution) or motor decision systems (e.g., response scaling).

### Behavioral experiment

Along with reproducing the center bias and serial dependence in heading perception from optic flow (e.g., [15-19]), the current experiment’s first contribution is to resolve the controversy that heading perception from optic flow shows either repulsive serial dependence [17] or attractive serial dependence [18] in different studies. Sun et al. [17] first examined the serial dependence in heading perception and found a repulsive serial dependence. Before their study, most studies consistently found attractive serial dependence in various features, such as orientation [23], color [41], and numerosity [28]. They proposed that their repulsive serial dependence was due to the dynamic feature of optic flow (also see the facial expression perception in [42]). However, Alais et al. [26] found an attractive serial dependence in motion direction perception, challenging the dynamic proposal. In particular, Xu et al. [18] found an attractive serial dependence in heading perception from optic flow. After comparing these studies, we found that the attractive serial dependence occurred when the stimulus values were uniformly distributed (e.g., [18, 26]), and the repulsive serial dependence occurred when the stimulus values followed a Gaussian distribution (e.g., [17]). Therefore, the distribution of stimulus values might affect serial dependence. Our current study well supported our hypothesis, confirming the effects of the stimulus distribution on serial dependence. Additionally, the current experiment was the first to demonstrate the effects of long-term experience on visual perception with experimental methods. As mentioned in the introduction, we defined long-term experience as the statistical regularity learned daily or in the long evolutionary process, which is different from the experience learned within a few days. Therefore, we cannot directly manipulate it with experimental methods. Previous studies have generally discussed its effects on visual perception through computational modeling (e.g., prior in the Bayesian model, [18, 43-46]). Levari and his collaborators found that when one feature’s proportion was reduced compared with its proportion a few minutes ago, participants tended to bias their perception more toward that feature, named the PICC effect [20, 21]. Inspired by their studies, we asked if the long-term experience affected our visual perception, then similar result trends could be observed by varying the proportions of features in the short-term experience to create different experience conflicts. Our results confirmed our expectations and demonstrated the effects of the long-term experience on our visual perception.

Moreover, the current finding of the effects of the conflict between short-and long-term experiences on heading estimation error led us to look carefully at the estimation errors of other visual features in previous studies. For example, huge studies have revealed a tilt or oblique illusion on orientation perception, meaning that participants bias their orientation estimates away from the oblique orientation [11-14]. Based on the environment’s statistics, Girshick et al. [10] proposed that the proportions of vertical and horizontal orientations were more than the oblique orientations (also see [9]), following an equation p(θ) 2-abs(sin(θ)) [44]. However, in tilt illusion studies, researchers generally selected oriented stimuli from uniform distributions, leading to the proportions of vertical and horizontal orientations in the short-term experience being less than that in the long-term experience. According to our current findings, participants may bias their orientation estimates toward the vertical and horizontal orientations, leading to a weakening of the oblique effect in these previous studies. That is, when the experience conflict disappears, the oblique effect may be stronger. Another example is visual speed perception. Previous studies have shown an underestimation of speed [47-49]. Weiss, Simoncelli, and Adelson [50] proposed that slower speeds are more likely to occur than faster ones (also see [38]). As in the orientation studies, the researchers selected their slow speeds from a uniform distribution [47, 48], which resulted in the proportion of slow speed in the short-term experience being significantly smaller than that in the long-term experience. As a result, the observed speed underestimation became significant. Hence, given the finding of the experience conflict effect on heading perception from optic flow, the question arises as to what participants’ actual estimates would be if we selected stimulus values in the experimental design according to the long-term experience. Meanwhile, our findings prompt us to carefully consider the selection of stimulus values in the experimental design because different distributions of stimulus values can generate different results.

### Bayesian inference modeling

The good predictions of our Bayesian models also supported the claim that heading perception from optic flow is consistent with the Bayesian inference theory, in which observers would rely more on the prior (i.e., long-term experience) to estimate their headings when the certainty of the stimuli’s sensory representation reduced [39]. Our Model 1 assumed that the high frequently present peripheral headings in the hill and flat distributions made the sensory neurons (e.g., MSTd, [51, 52]) fatigued, leading to a decrease in the certainty of the sensory representation (i.e., an increase in the standard deviation of likelihood distributions). As a result, the model well captured the increasing trend of center bias when the distribution became hill and flat from the peak, consistent with participants’ performances.

Additionally, previous studies have proposed that serial dependence in visual perception also follows the Bayesian inference theory [18, 33, 53]. Xu et al. [18] first developed a Bayesian model and captured the attractive serial dependence in heading perception from optic flow. In their model, the heading estimate resulted from the optimal combination of the prior distribution and the likelihood distributions of previous and current headings. Note that their Bayesian model only predicts the attractive serial dependence but does not work for the repulsive serial dependence. In the current study, we used the Bayesian model to simulate participants’ estimates and analyzed the serial dependence in these simulated data. We captured both repulsive and attractive serial dependence well. Therefore, our model further supported the Bayesian inference process of serial dependence.

Moreover, our models explained the potential mechanisms underlying the effects of the experience conflict on heading perception from two perspectives. One possibility is that the experience conflict increased the fatigue of sensory systems, making the likelihood distributions uncertain (red arrows in Figure 6). Hence, participants relied more on the prior distribution and previously present heading, showing stronger center bias and attractive serial dependence, consistent with participants’ performance (Figure 2a). Another possibility is that the experience conflict changed participants’ motor systems. Specifically, when there were more peripheral headings in the short-term experiences, participants might avoid reporting peripheral headings and shrink their response ranges. Therefore, our Model 2 worked like the classical Bayesian model (gray shaded area in Figure 6) but multiplied an extra response scaling factor (Equation 7) with the decoded estimates. Model 2 also captured the participants’ performances, suggesting that the experience conflict affected heading perception by influencing motor decision systems.

Therefore, we developed two Bayesian inference models to explain our findings. Some researchers could argue that why not use classical Bayesian inference model (gray shaded area in Figure 6) to optimally combine the stimulus distribution, prior distribution and the likelihood distribution. According to the Bayesian inference theory (Equation 4) or ideal observer model [54, 55], our final estimate can be directly given by:

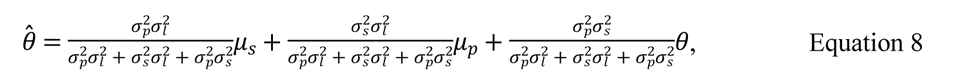

in which, *σ*_*p*_, *σ*_*s*_ and *σ*_*l*_ represent the standard deviations of the prior distribution, stimulus distribution and likelihood distribution. *μ*_*p*_, *μ*_*μ*_ and *θ* represent the means of the three distributions. Among them, *μ*_*p*_ and *μ*_*s*_ were zero, *θ* corresponds to the actual headings.

Therefore, Equation 8 can be simplified into:

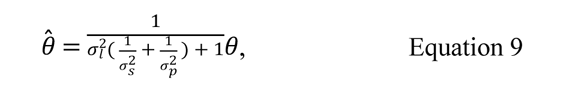

We know that the size order of *θ*_s_S is peak < hill < flat distributions. Hence, with Equation 9, *θ* in the peak distribution is the smallest, showing strongest center bias; in contrast, *θ* in the flat distribution is the largest, showing weakest center bias. That is, the predications of the classical Bayesian inference model are opposite to participants’ performance.

Lastly, the prior and likelihood distributions were independent in our model. Wei and Stocker [44-46] developed a Bayesian efficient-coding model in which the prior constraints our likelihood distribution and the decoding process. In addition, they argued that noise affecting visual perception could be divided into two types: sensory vs. physical stimulus noises. They found that increasing the physical noise made the features more underestimated (e.g., an increase in center bias in heading estimation) [44, 45]. However, optic flow stimuli were the same across different distributions in our current study, so the physical noises were constant. Hence, Wei’s Bayesian efficient coding model could not explain our findings.

### Future studies

The current study proposed that the effects of the conflict between short-and long-term experiences could be due to the changing of the certainty of sensory systems’ representations and response decision systems. However, the validity of the above conclusions is still an open question because the above conclusions were made by developing computational models. Hence, one future direction is to examine the validity of our findings. Physiological and brain imaging studies have demonstrated that several cortical areas in our brains are involved in the representation of heading perception from optic flow, such as MT/V5, MST, V3b/KO (see [56] for review), VIP and CSv [57-59]. In addition, the frontal and parietal cortical circuits are responsible for motor decision-making (see [60] for review). If the experience conflict affected heading perception by modulating sensory systems’ representation, then these cortical areas’ activities of sensory systems would be different in different experience conflict conditions. Similarly, if the experience conflict works by modulating motor decision systems, then these cortical areas’ activities of motor systems would be different in different experience conflict conditions. Therefore, in the future, we can consider adopting brain imaging (e.g., fMRI) or electrophysiological (e.g., EEG) techniques to record these cortical areas’ activities in different experience conflict conditions to answer the role of sensory and motor systems in the effect of experience conflict.

Moreover, Levari’s PICC effect is also applicable in the facial threat and research ethical judgment [20], suggesting that Levari’s PICC effect covers physical features (e.g., color) and some complex social features. The stimuli in the current study are optic flow, a middle-level physical feature. The natural world has many physical features, such as low-level features: colors, orientations; and high-level features: facial expressions and emotions. Hence, whether our findings can be extended to these features remains unclear.

As mentioned above, the long-term experience investigated in the current study corresponds to be the prior distribution in the Bayesian inference model [18, 43-46]. Previous studies built the prior distribution based on natural [10] or learned [2] stimulus statistic. The current study provides a new way to deduce the prior distribution. Specifically, if we propose that one feature estimation follows the Bayesian inference rule, we can set several distributions of features and collect participants’ estimates in different conditions. By analyzing their data trends, we can develop the prior distribution. Therefore, the current experiment provided a practicable experimental method or idea for the prior development.

To sum up, the current study may set several avenues for future studies.

## Conclusions

In conclusion, the current study revealed that the conflict between short-and long-term experiences affected our perceptual error and serial dependence in visual perception, and showed that the effects were consistent with a Bayesian inference account. We proposed that the experience conflict affected visual perception by modulating our sensory or motor decision systems. Our current study not only improves our understanding of the processing mechanisms underlying visual perception, but also provides some new ideas or inspiration for researchers on research questions and experimental methods.

## Author Contributions

QS: Conceptualization, Data Curation, Formal Analysis, Funding Acquisition, Methodology, Project Administration, Resources, Software, Supervision, Validation, Visualization, Writing-Original Draft Preparation, Writing-Review & Editing. JYW and XMG: Investigation, Validation, Writing-Original Draft Preparation, Writing-Review & Editing.

## Acknowledgement

The study was supported by National Natural Science Foundation of China, China (No. 32200842) to Qi Sun.

## Supporting information captions

Participants’ data in the current study can be downloaded from the link of OSF: https://osf.io/zr8jp/

